# Immune disease dialogue of chemokine-based cell communications as revealed by single-cell RNA sequencing meta-analysis

**DOI:** 10.1101/2024.07.17.603936

**Authors:** Mouly F. Rahman, Andre H. Kurlovs, Munender Vodnala, Elamaran Meibalan, Terry K. Means, Nima Nouri, Emanuele de Rinaldis, Virginia Savova

## Abstract

Immune-mediated diseases are characterized by aberrant immune responses, posing significant challenges to global health. In both inflammatory and autoimmune diseases, dysregulated immune reactions mediated by tissue-residing immune and non-immune cells precipitate chronic inflammation and tissue damage that is amplified by peripheral immune cell extravasation into the tissue. Chemokine receptors are pivotal in orchestrating immune cell migration, yet deciphering the signaling code across cell types, diseases and tissues remains an open challenge. To delineate disease-specific cell-cell communications involved in immune cell migration, we conducted a meta-analysis of publicly available single-cell RNA sequencing (scRNA-seq) data across diverse immune diseases and tissues. Our comprehensive analysis spanned multiple immune disorders affecting major organs: atopic dermatitis and psoriasis (skin), chronic obstructive pulmonary disease and idiopathic pulmonary fibrosis (lung), ulcerative colitis (colon), IgA nephropathy and lupus nephritis (kidney). By interrogating ligand-receptor (L-R) interactions, alterations in cell proportions, and differential gene expression, we unveiled intricate disease-specific and common immune cell chemoattraction and extravasation patterns. Our findings delineate disease-specific L-R networks and shed light on shared immune responses across tissues and diseases. Insights gleaned from this analysis hold promise for the development of targeted therapeutics aimed at modulating immune cell migration to mitigate inflammation and tissue damage. This nuanced understanding of immune cell dynamics at the single-cell resolution opens avenues for precision medicine in immune disease management.

## 1. INTRODUCTION

Immune diseases are complex and often devastating disorders in which the immune system goes rogue. In both inflammatory and autoimmune diseases such as atopic dermatitis (AD) and ulcerative colitis (UC), the immune response to allergens (Roesner et al., 2016) or autoimmune activity (Zhonghui and Claudio, 2004; Gajendran et al., 2019) involves interactions between tissue-residing immune cells and non-immune endothelial, epithelial and fibroblast cells. These interactions result in the secretion of inflammatory mediators, including chemokines, which drive immune cell chemotaxis (Banks et al., 2003; Gros et al., 2009; Shachar and Karin, 2013). This inflammatory cascade may be amplified by neighboring tissue-resident immune cells which further facilitate the arrival of other immune cells to the site of inflammation (John and Abraham, 2013; Linthout et al., 2014). Additionally, blood vessel endothelium upregulates integrins, cadherins and selectins, adhesive proteins that drive peripheral immune cell extravasation (Luster et al., 2005). Utilizing a diverse vocabulary of ligand-receptor (L-R) signals, these immune cells invade the tissue and engage in complex cellular crosstalk, leading to localized chronic inflammation, tissue damage, and even impaired organ function (Chen et al., 2017; Tsokos, 2020).

Chemokine receptors have been a focus of therapeutic targeting due to their crucial role in directing immune cells to an inflammatory environment (Lai and Mueller, 2021). However, these attempts have been largely unsuccessful, possibly due to the complex nature of chemokine receptors, which can bind multiple chemokines (Solari et al., 2015; Lai and Mueller, 2021). Furthermore, the specific effects of chemokine L-R signals in relation to diseases, tissues, or cell types remains unclear (Stone, 2017), especially in the context of human tissue. The emergence of single-cell RNA sequencing (scRNA-seq) enables the analysis of gene expression at the individual cell level within heterogeneous tissues, offering experimental evidence for potential L-R pairs and cell-cell communication in a high-throughput manner (Heumos et al., 2023). Additionally, it allows for the identification of cell-type-specific gene expression changes, aligning with the understanding that human diseases often manifest in a tissue and cell-specific manner (Hekselman and Yeger-Lotem, 2020). As technology improves, scRNA-seq datasets are gradually revealing altered cellular phenotypes associated with various immune diseases affecting millions globally (Conrad et al., 2023), including AD, psoriasis (PSO), chronic obstructive pulmonary disease (COPD), idiopathic pulmonary fibrosis (IPF), UC, immunoglobulin A nephropathy (IgAN) and lupus nephritis (LN). However, a meta-analysis of scRNA-seq datasets to find overlapping and tissue/disease-specific cell-cell communications is currently unavailable.

Utilizing information from publicly available scRNA-seq datasets profiling a diverse set of patient samples, we aimed to evaluate common and disease-specific L-R interactions across various diseases and tissues. To enhance the reliability of our findings, we conducted a meta-analysis that spanned multiple diseases and tissues to minimize the impact of batch, experimental, or patient-specific effects. The objective was to provide valuable insights supporting the clinical development of disease and tissue-specific therapeutics. Specifically, we performed a ligand-receptor analysis of scRNA-seq data with the focus on chemokine genes to identify disease and/or tissue-specific patterns. Considering that the receiver cells involved in chemokine L-R interactions could be either resident or non-resident peripheral immune cells that have extravasated into the tissue, we conducted additional ligand-receptor analyses focused on immune cell extravasation genes, assessing interactions between endothelial cells and immune cells to infer whether peripheral immune cells had indeed infiltrated the tissue. Furthermore, we examined changes in cell proportions in disease tissues compared to their healthy controls, as well as differentially expressed gene (DEG) analysis encompassing chemokines, immune cell extravasation, cell proliferation, and activation markers, aiming to understand whether increased cell proportion, activation, or gene expression may be related to enhanced L-R interactions for those cells. As this comprehensive approach allowed us to identify potentially relevant cell-cell L-R interactions that have not been investigated in depth in past research, we proceeded to experimentally validate these findings. This validation involved subjecting the purportedly migrated cell to chemotaxis assays, wherein the candidate ligand served as the chemoattractant. In certain instances, we took an additional step by implementing a knockout of the candidate receptor to assess whether this would impede chemoattraction as hypothesized.

## 2. METHODS

### 2.1 Single-cell RNA sequencing dataset selection and processing

We selected diseases based on the availability of public scRNA-seq datasets across skin, lung, colon and kidney healthy control tissue, and their respective diseases (encompassing AD, PSO, COPD, IPF, UC, IgAN, LN) and at least two samples per group. We collected and analyzed the following 15 scRNA-seq datasets in our analysis: 1) E-MTAB-8142 (Reynolds et al., 2021) for healthy skin, AD lesional skin, and AD nonlesional skin, PSO lesional skin and PSO nonlesional skin, 2) GSE147424 (He et al., 2020) for healthy skin, AD lesional skin, and AD nonlesional skin, 3) GSE153760 (Rindler et al., 2021) for healthy skin and AD lesional skin, 4) GSE173706 (Merleev et al., 2022) for healthy skin, PSO lesional skin, and PSO nonlesional skin, 5) GSE220116 (Kim et al., 2023) for healthy skin and PSO lesional skin, 6) EGAS00001004344 (Travaglini et al., 2020) for healthy lung, 7) GSE136831 (Adams et al., 2020) for healthy lung, COPD lung and IPF lung, 8) GSE171541 (Huang et al., 2022) for healthy lung and IPF lung, 9) GSE122960 (Reyfman et al., 2018) and 10) GSE135893 (Habermann et al., 2020) for healthy lung and IPF lung, 11) SCP259 (Smillie et al., 2019) and 12) GSE116222 (Parikh et al., 2019), and 13) GSE231993 (Du et al., 2023) for healthy colon, UC inflamed colon, and UC uninflamed colon, 14) GSA: HRA000342 (Zheng et al., 2020) for healthy kidney and IgAN kidney, and 15) LN_Kidney_AMP (from the Accelerating Medicines Partnership (AMP) SLE phase 2 consortium) for healthy kidney and LN kidney (Arazi et al., 2019; Hoover et al., 2023; Horisberger et al., 2024; Izmirly et al., 2024).

We processed the publicly deposited count matrices for these datasets using CellBridge (version 1.0.0; (Nouri et al., 2023)), an automated, Docker-based scRNA-seq data processing pipeline developed by our research group, with empirical filtering parameters for each dataset (specified in **Supplemental Table 2**) and specific pipeline steps described hereafter. Doublet removal was performed using the Scrublet Python package (version 0.2.3; (Wolock et al., 2019)). The Seurat R package (version 5.0.1; (Hao et al., 2024)) was used to perform initial processing (“NormalizeData”, “FindVariableFeatures”, “ScaleData”, “RunPCA” with default parameters). To improve our cell type annotation (which relies on dimensionality reduction for smoothing), sample-based batch correction of the PCA was performed using the Harmony R package (version 1.0; (Korsunsky et al., 2019)), followed by the nearest-neighbor graph construction using “FindNeighbors” with 30 dimensions (with default settings except “dims=1:30”), subsequent cluster identification with “FindClusters” (with default settings except “resolution=0.7”), and the Uniform Manifold Approximation and Projection (UMAP) dimensionality reduction with “RunUMAP” on 30 dimensions (with default settings except “dims=1:30”). Cell type annotation was performed by the SignacX R package (version 2.2.5; (Chamberlain et al., 2023)), also produced from our group, that is a classifier designed to predict cellular phenotypes in scRNA-seq data from multiple tissues. Cell classifications from SignacX include: memory B cells (“B.memory” in SignacX nomenclature), naïve B cells (“B.naive”), dendritic cells (“DC”), endothelial, epithelial, fibroblasts, macrophages, classical monocytes (“Mon.Classical”), non-classical monocytes (“Mon.NonClassical”), neutrophils, natural killer cells (“NK”), plasma cells (“Plasma.cells”), CD4+ naïve T cells (“T.CD4.naive”), CD4+ memory T cells (“T.CD4.memory”), CD8+ naïve T cells (“T.CD8.naive”), central memory CD8+ T cells (“T.CD8.cm”), CD8+ effector memory T cells (“T.CD8.em”), and regulatory T cells (“T.regs”).

### 2.2 Cell-cell communication analysis

To identify potential ligand-receptor interactions taking place between cells, we used CellphoneDB version 5.0, which contains a curated repository of ligand–receptor interactions and a permutation-based statistical framework for inferring enriched interactions between cell types from scRNA-seq data (Efremova et al., 2020). Count matrices and metadata files extracted from the Seurat RDS provided by CellBridge were processed using the default parameters of the statistical method of CellphoneDB v.5.0 (https://github.com/ventolab/CellphoneDB) in Python (version 3.7; Python Software Foundation). We filtered the CellphoneDB statistically significant means for interactions where either the ligand or the receptor is a chemokine gene (i.e. HGNC gene symbol starting with CC or CX) and from a list of genes known to be involved in immune cell extravasation (listed in **Supplemental Table 1;** (DeGrendele et al., 1997; Johnson-Léger et al., 2002; Ludwig et al., 2005; Farkas et al., 2006; Wegmann et al., 2006; Engelhardt, 2008; Kumar et al., 2012; Azcutia et al., 2013; Rocha et al., 2014; Baeyens et al., 2015; Glatigny et al., 2015; Zhang et al., 2016; Bros et al., 2019; Bui et al., 2020; Arif et al., 2021; Amersfoort et al., 2022; Czubak-Prowizor et al., 2022; Ma et al., 2023; Xu et al., 2023)). For the extravasation-based output, the sender cell (i.e. the cell expressing the ligand) was filtered to Endothelial cells, as this is the cell type involved in mediating immune cell extravasation through blood vessel endothelium (Luster et al., 2005).

### 2.3 Cell proportion comparisons

We conducted pairwise (disease vs healthy) comparisons of cell type proportions using the Brunner-Munzel test in the brunnermunzel R package (version 2.0) on each dataset. Analysis was done with the minimum number of subjects per group to hold a comparison (per cell state) set to two, and the minimum number of total cells per subject set to 200. Results with Benjamini-Hochberg-corrected p-values <0.05 were considered statistically significant.

### 2.4 Differentially expressed gene analysis

We conducted pairwise (disease vs healthy) DEG analysis using the zero-inflated negative binomial model (zlm) in the model-based analysis of single-cell transcriptomics (MAST) R package (version 1.26.0; (Finak et al., 2015)) on each cell type in each dataset with sample covaried. For a gene to be considered, it had to be expressed in at least 25% of cells in a given cell type and have an absolute log2-fold change of at least 0.25. Genes with Benjamini-Hochberg-corrected false discovery rate (FDR) of <0.05 were further filtered for our target genes, which included chemokine genes (i.e. HGNC gene symbol starting with CC or CX), immune cell extravasation, proliferation, and activation genes (listed in **Supplemental Table 1** as indicated earlier).

### 2.5 Meta-analysis

After conducting each analysis (cell-cell communication, cell proportion, DEG) per dataset, RStudio (RStudio Team, version 4.2) was used to extract common statistically significant findings across datasets, within each disease group in the case of cell proportion, DEG and cell-cell communication analyses, as well as the corresponding healthy groups (healthy skin, healthy lung, healthy colon, healthy kidney) in the case of cell-cell communication analyses.

For cell-cell communication, interactions that were statistically significant (CellphoneDB’s permutation test, p < 0.05) were selected for further analysis. We analyzed whether an interaction was multi-tissue (i.e. across multiple (healthy or disease) tissues), or disease-enriched (i.e. specific to diseased tissues only), which could then be further subdivided according to specificity, from occurring in multiple diseases across tissues, to occurring only in diseases of specific tissues (e.g., skin), to those only occurring in a particular disease. Visualizations were performed in RStudio (RStudio Team, version 4.2) using the ggplot2, ggsankey, and circlize libraries (Gu et al., 2014).

### 2.6 Transwell chemotaxis assay

To validate interactions from the cell-cell communication analyses, we chose two intriguing findings that had been unexplored in the literature in terms of chemotaxis assays: chemokine (C-X-C motif) ligand 2 (CXCL2) signaling (from various immune cells) to T.CD8.em cells (via dipeptidyl peptidase-4 (DPP4)) and retinoic acid receptor responder protein 2 (RARRES2) signaling to classical and nonclassical monocytes. First, we measured chemotaxis of pan CD8+ T cells (Charles River, cat. PB08NC-2) and pan monocytes (Charles River, cat. PB14-16NC-1) from 2-3 donors towards CXCL2 (R&D Systems, cat. 276-GB-010) and RARRES2 (R&D Systems, cat. 224-CM-025), respectively, using a transmigration chamber assay with 5 μm pore size transwell inserts (Corning, cat. CLS3421) according to the manufacturer’s instructions. Briefly, the transwell inserts were placed into 24-well plates containing 500 ul of Roswell Park Memorial Institute (RPMI) 1640 media (Gibco, cat. 22400-089) with 1% heat-inactivated fetal bovine serum (FBS; Gibco, cat. A38400-01, lot. 2717806RP) alone (the medium only negative control condition), medium with a positive control, or medium with each target ligand (at multiple dilutions). 2-4 wells were used per condition. For the CD8+ T cell experiments, chemokine (C-X-C motif) ligand 12 (CXCL12, 10 ng/ml; R&D Systems, cat. 350-NS-010) was used as the positive control. For the monocyte experiments, two positive controls were used; 5% (v/v) cobra venom activated human complement serum (CAS; Complement Technology Inc, cat. NC1769554), as well as CC motif chemokine ligand 8 (CCL8; R&D Systems, cat. 281-CP-010). Cells were quickly thawed in a 37 C water bath, spun down at 300g for 10 min and allowed to rest for 1 hour in an incubator (37 C, 5% CO2) in RPMI 1640 with 10% FBS, prior to being spun down at 300g for 10 min, counted with the CellacaMX (Nexcelom Bioscience), and then resuspended in 1% FBS media and loaded at a 100 ul volume, at 200,000 cells/transwell insert for the CD8+ T cell experiments and 250,000 cells/transwell insert for the monocyte experiments. Transwell plates were then incubated for 3h to permit chemotaxis. Cell migration was quantified using CellTiter-Glo (Promega, cat. G9241) relative luminescence (RLU) measured with a plate reader (EnVision, 2104-0010). To find significant changes in cell migration of each concentration of tested ligand compared to the medium control, student t-tests were performed on R studio (version 4.2). For each donor, migration index (MI) was calculated using the following formula and used for plotting.

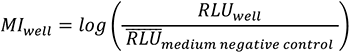

### 2.7 Flow cytometry and CRISPR knock-out

After conducting the initial chemotaxis experiments, we investigated whether knock-out of the receptor DPP4, identified through cell-cell communication analyses in mediating CXCL2 binding in CD8+ effector memory T cells, inhibits chemotaxis, and whether the expression of DPP4 changes post-chemotaxis.

Firstly, on day 0, a subset of CD8+ T memory cells (cluster of differentiation 45 restricted (CD45RO+); STEMCELL Technologies, cat. 200-0168) from two donors underwent the chemotaxis assay as described in the previous section. Transwell-migrated cell solutions (two sets of two wells pooled, per condition) were used for flow cytometry analysis for the receptor genes of interest. These cells along with compensation beads were placed in a 96-well clear round bottom plate (Corning, cat. 3799), washed twice by spinning down at 350g for 5 min and resuspended with 1X PBS (Gibco, cat. 10010049), stained in the dark for 15 min at room temperature (RT) with Zombie NIR Fixable viability dye (Biolegend, cat. 423105), washed twice with 1X PBS, stained in the dark for 30 min at 4C with an extracellular panel of antibodies at 1:100 dilution (Pacific blue CD8 (cat. 344717), BV570 CD45RO (cat. 304225), BV785 CD25 (cat. 302637), FITC DPP4 (cat. 302638), PE CXC receptor 4 (CXCR4, cat. 306505), PE/Dazzle 594 CD62 L-selectin (CD62L; cat. 304841), APC CXCR2 (cat. 320710); all anti-human and Biolegend) with Fc block (Biolegend, cat. 422302), washed twice with cell staining buffer (Biolegend, cat. 420201), permeabilized and fixed (using Fixation/Permeabilization Concentrate, ThermoFisher, cat. 00-5123-43 and Fixation/Permeabilization Diluent, ThermoFisher, cat. 00-5223-56) for 30 min at RT, washed twice with perm/wash buffer (ThermoFisher, cat. 00-8333-56), stained in the dark again with the CXCR4, CXCR2 and DPP4 antibodies and Fc block for intracellular staining, washed twice with cell staining buffer, resuspended in 40 ul cell staining buffer, and quantified with the iQue3 (Sartorius) at a sip time of 37s.

Analysis was conducted on OMIQ (Dotmatics); unfiltered data was gated for cells, followed by live/dead gating, singlets, and then fluorescence minus one (FMO)-based gating of CD8+ cells, followed by the target proteins (CXCR4, CXCR2, DPP4, CD25). Percentage of cells positive for each marker (receptors including CXCR4, CXCR2, DPP4 and activation marker CD25) and their fluorescence intensity was compared to the medium negative control with student t-tests in R Studio (version 4.2).

The remaining CD8+ memory T cells not used for the chemotaxis assay were incubated in a T25 flask with ImmunoCult-XF T cell expansion medium (ICEM; STEMCELL Technologies, cat. 10981) and 50 IU/mL of IL-2 (STEMCELL Technologies, cat. 78220) at 1 million cells/mL. On day 1, cells were activated with 25 ul/mL CD3/28 activator (STEMCELL technologies, cat. 10971). On day 4, cells were divided amongst the following knock-out groups: wild-type (WT), “zap-only”, DPP4 knock-out (KO), and non-target guide control (NTC). The DPP4 KO and NTC groups were washed with 1X PBS and resuspended with P3 primary nucleofection solution (Lonza, cat. V4XP3012) containing ribonucleoprotein (RNP) complexes of 20 pmol cas9 nuclease protein (Horizon Discovery, cat. CAS12205) and 200 pmol guide RNA (gRNA) per 1 million cells, targeting either DPP4 (Horizon Discovery, cat. SQ-004181-01-002) or the NTC locus (Synthego, CRISPRevolution sgRNA EZ kit). The RNPs were assembled by 37 C incubation for 10 minutes, followed by RT incubation for 5 min. The cell-RNP mixture was nucleofected using an electroporator (4D-Nucleofector X Unit, Lonza) according to the manufacturer’s instructions, and then incubated with ICEM in a 12-well plate. The “zap-only” group underwent the same procedure, but without any introductions to RNPs, and the WT group was simply washed with 1X PBS, resuspended in ICEM and transferred to the 12-well plate. On day 11, 100,000 cells per group and per donor underwent the flow cytometry protocol outlined earlier to verify whether DPP4 was KO’d in the DPP4 KO group and still expressed amongst the control groups (WT, zap-only and NTC). On day 12, WT, DPP4 KO and NTC groups were used in the CXCL2 chemotaxis assay (with medium and CXCL12 controls) for chemotaxis and flow cytometry quantification as described earlier.

## 3. RESULTS

In this study, we leveraged 15 public scRNA-seq human immune disease datasets to identify tissue- and disease-specific, as well as common, chemokine and extravasation ligand-receptor cell communications that could be contributing to the recruitment of immune cells to the tissues. We started by using CellphoneDB to identify statistically significant cell-cell communications, followed by DEG analysis to determine which of the genes involved in cell-cell communication were differentially expressed in diseased vs healthy tissue, as well as cell proportion analysis that shed light on the interplay between cell communication and cell abundance. Finally, based on our findings from the public datasets, we conducted experimental validation of two previously understudied interactions using chemotaxis assays in the laboratory.

### 3.1 Datasets and meta-analysis

We curated 15 datasets representing the following: 1) three datasets of AD skin (lesional in all and non-lesional in two of them), 2) three datasets of PSO skin (lesional in all and non-lesional in two of them), 3) two datasets of IPF lung (one of which also contained COPD) and another lung dataset containing COPD, 4) three datasets of UC colon (inflamed and non-inflamed), and 5) two kidney datasets, one of IgAN, and another one of LN. All datasets contained healthy controls as well (**Figure 1a**).

**Figure 1.**
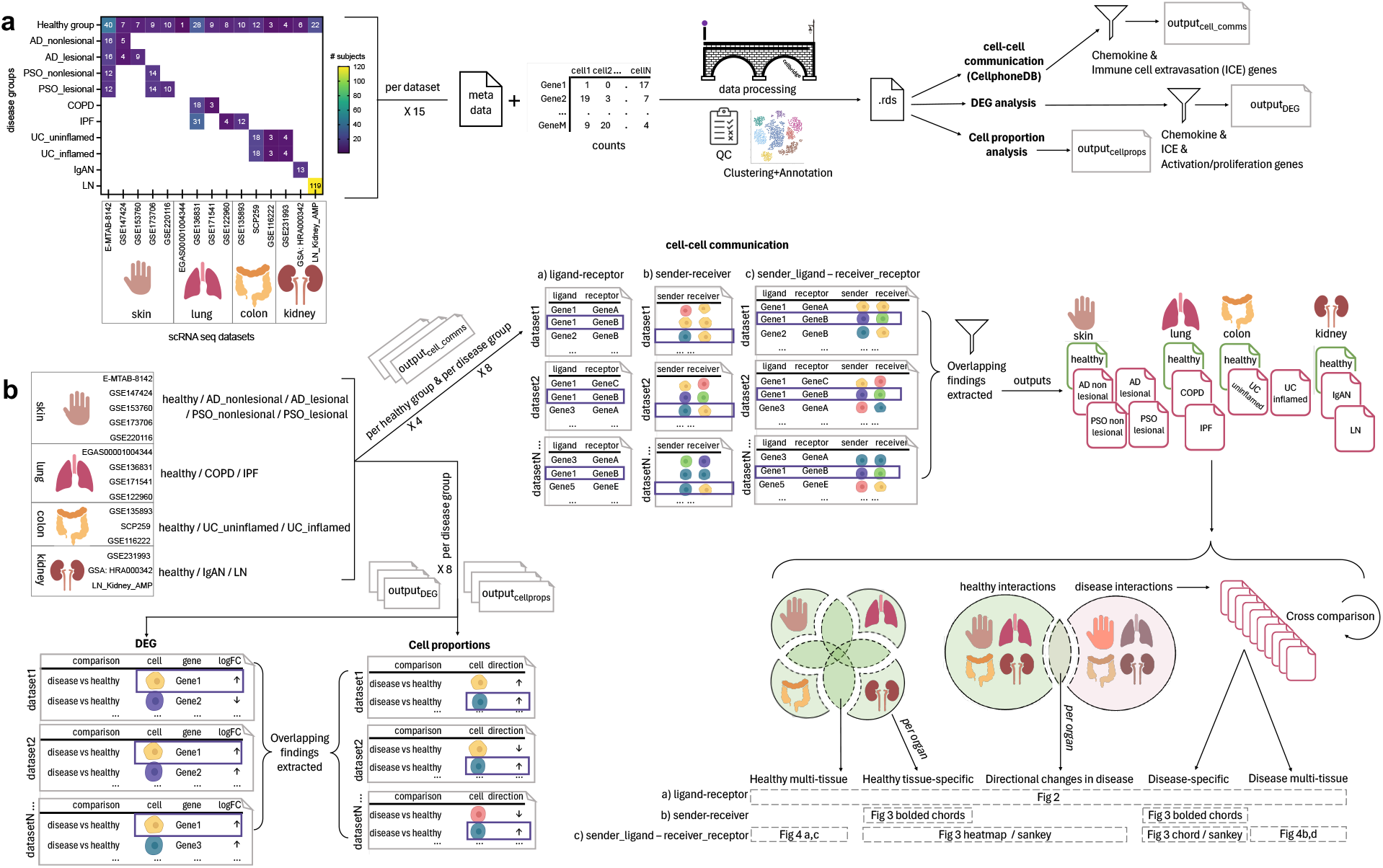
Overview of the single cell (sc)-RNA-seq datasets selected and how they were processed and analyzed for the meta-analysis of chemokine and extravasation-based cell-cell communication. **a)** 15 scRNA-seq datasets were selected, encompassing the following 14 groups: healthy skin, atopic dermatitis (AD) nonlesional skin, AD lesional skin, psoriasis (PSO) nonlesional skin, PSO lesional skin, healthy lung, chronic obstructive pulmonary disease (COPD) lung, idiopathic pulmonary fibrosis (IPF) lung, healthy colon, ulcerative colitis (UC) uninflamed colon, UC inflamed colon, healthy kidney, IgA nephropathy (IgAN) kidney, lupus nephritis (LN) kidney. The counts and metadata files for each dataset were processed using the CellBridge pipeline. The Seurat RDS output was used for the down-stream analyses: cell-cell communication using CellphoneDB, differentially expressed gene (DEG) analysis (disease vs healthy), and cell proportion (disease vs healthy) analysis. Cell-cell communication results were filtered for interactions involving chemokine and immune cell extravasation genes, and DEG results were filtered for chemokine, immune cell extravasation, immune cell activation and proliferation genes. **b)** Per a set of datasets belonging to a particular group, common findings were extracted for each of the three types of analyses (cell-cell communication, DEG, cell proportions). For the cell-cell communication results: 1) healthy-tissue specific findings were extracted by comparing each tissue to each other (e.g. to extract interactions only occurring in healthy kidney), 2) all healthy interactions were compared to all disease interactions to extract those only occurring in disease, and then 3) those disease interactions were compared against each other to extract interactions that were specific to a particular disease group (e.g. only in COPD). These comparisons resulted in findings that were: healthy multi-tissue, healthy tissue specific, directional changes in disease (i.e. healthy interactions that occurred in disease, but with an increased/decreased interaction value, hence the ‘directional’ change), disease-specific and disease multi-tissue.

After conducting each form of analysis (cell-cell communication via CellphoneDB, cell proportion, DEGs) per dataset, we extracted findings that were present in all datasets of a particular disease group (AD nonlesional skin, AD lesional skin, PSO nonlesional skin, PSO lesional skin, COPD lung, IPF lung, UC uninflamed colon, UC inflamed colon, IgAN kidney, LN kidney) in the case of cell-proportion, DEG and cell-cell communication analyses, as well as healthy group (healthy skin, healthy lung, healthy colon, healthy kidney) in the case of cell-cell communication analyses (**Figure 1b**). Statistical results for the cell proportion data are reported in **Supplemental Table 3** and for the DEG data reported in **Supplemental Table 4.**

For cell-cell communication analysis, after extracting findings that were present in all datasets of a particular disease group and healthy group, the next process involved four steps: 1) comparing CellphoneDB findings from each disease tissue with those from all healthy tissues to extract ‘disease-enriched’ interactions, 2) further comparing these ‘disease-enriched’ findings against each other to extract disease-specific interactions (e.g. interactions only present in COPD lung but in no other healthy or disease group), 3) comparing findings from each healthy tissue against those from every other healthy tissue to extract healthy-tissue-specific interactions (e.g. interactions only present in healthy lung, but not in healthy skin, colon or kidney), and 4) extracting healthy interactions that overlapped with disease tissue interactions. These CellphoneDB cell-cell communications and their interaction values found in healthy tissues are reported in **Supplemental Table 5**, interactions that are disease-enriched are reported in **Supplemental Table 6**, and healthy tissue interactions that also occurred in disease tissue are reported in **Supplemental Table 7**.

### 3.2 Ligand-receptor pairs identified by the cell-cell communication analysis

Overall, we identified 39 unique chemokine-based L-R pairs by cell-cell communication analysis (**Figure 2a**). The majority of these interactions (28 or 72%) were “disease-enriched”, i.e., they did not occur in any healthy tissue. Of these disease-enriched L-R pairs, 11 (28%) were specific to a particular disease (referred to as “disease-specific”), 12 (31%) were specific to a particular organ but co-occurring in multiple diseases of that organ (referred to as “disease organ-specific”), five (13%) occurred in various diseases (referred to as “disease multi-tissue”). Interestingly, among the disease multi-tissue L-R pairs, the sender and receiver cell types implicated in their expression were entirely distinct depending on the diseased organ type. For example, the CCL3L1 --> CCR1 interaction was found in IgAN kidney, LN kidney and UC uninflamed colon; however, in UC uninflamed colon, the CCL3L1 --> CCR1 interaction occurred between Neutrophils --> Macrophages, whereas in IgAN or LN kidney, other cell types were involved in this communication.

**Figure 2.**
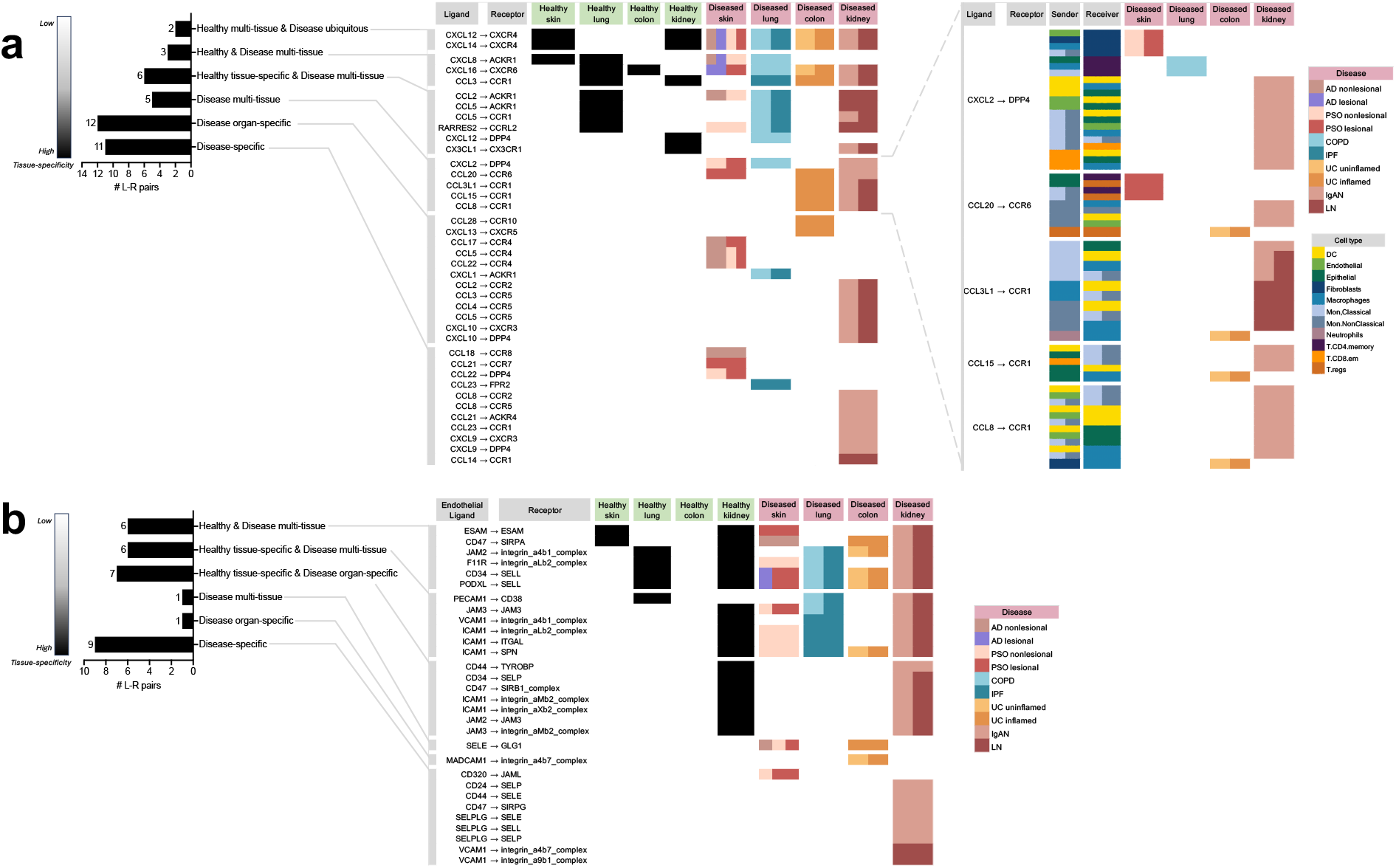
All of the unique chemokine and immune cell extravasation-based ligand-receptor pairs extracted from the meta-analysis of cell-cell communication predictions based on CellphoneDB. **a)** The chemokine-based ligand-receptor pairs that took place in: multiple healthy tissues and all diseases (“Healthy multi-tissue & Disease ubiquitous”), multiple healthy and disease tissues (“Healthy & Disease multi-tissue”), a specific healthy tissue along with multiple disease tissues (“Healthy tissue-specific & Disease multi-tissue”), only disease tissue and not in any healthy tissue, but not specific to a particular diseased tissue (“Disease multi-tissue”), a specific diseased organ (“Disease organ-specific”), a particular disease (“Disease-specific”). For “Disease multi-tissue”, sender-receiver cell type-pairs were completely unshared across disease organ types (shown in the blow-out). **b)** The immune cell extravasation-based Endothelial_ligand – receptor pairs that took place in the various sets mentioned above. *AD* atopic dermatitis, COPD chronic obstructive pulmonary disease, *IgAN* IgA nephropathy, *IPF* idiopathic pulmonary fibrosis, LN lupus nephritis, *PSO* psoriasis, *UC* ulcerative colitis.

Unlike the disease-enriched interactions, there were no L-R pairs that were “healthy-enriched”, i.e. only found in healthy tissues; of the remaining 11 (28%) interactions, all occurred in both disease and healthy tissue. Of those, six (16%) were specific to a particular healthy tissue, while also occurring in various diseases (referred to as “healthy tissue-specific & disease multi-tissue”), three (8%) were occurring in varieties of both healthy and disease tissues (referred to as “healthy & disease multi-tissue”), and two (5%) were occurring in various healthy tissues and ubiquitously occurring in all the diseases examined in the study (referred to as “healthy multi-tissue & disease ubiquitous”).

There were 30 unique extravasation-based L-R pairs identified by cell-cell communication analysis (**Figure 2b**). Unlike among the chemokine-based L-R pairs, a minority (33%) of extravasation- based L-R pairs were “disease-enriched’; of these interactions, nine (30%) were disease-specific, one (3%) was “disease organ-specific”, and one (3%) was in the “disease multi-tissue” set. The only “disease multi-tissue” L-R pair was SELE --> GLG1, which occurred in inflamed UC colon and nonlesional AD and PSO skin.

As with the chemokine-based L-R pairs, none of the extravasation-based L-R pairs were found exclusively in healthy tissue. However, contrary to the chemokine-based L-R pairs, the majority of the extravasation-based interactions (20; 67%) occurred in both healthy and disease tissue. Of the 20, seven (23%) were specific to a particular healthy tissue and specific to a diseased organ (referred to as “healthy tissue-specific & disease organ-specific”; interestingly, all of them were specific to the kidney), six (20%) were specific to a particular healthy tissue, while also occurring in various diseases (referred to as “healthy-tissue specific & disease multi-tissue”), and six (20%) were occurring in varieties of both healthy and disease tissues (referred to as “healthy & disease multi-tissue”). Of note, none of the extravasation- based L-R pairs were healthy multi-tissue & disease ubiquitous, as found in the chemokine-based L-R pairs.

### 3.3 Sender_ligand --> receiver_receptor pairs identified by the cell-cell communication analysis

Taking into account all four components of cell-cell communication — the sender cell, its ligand, the receiver cell, and its receptor — we proceeded to examine interactions specific to each of the healthy tissues studied (skin, lung, colon, and kidney) and their corresponding diseases. Intriguingly, in each of the healthy tissues (lung, skin, colon and kidney), sender-receiver pairs were unique to each tissue type regardless of the chemokine-based ligands and receptors. For example, in healthy lung, there was a Macrophage --> Endothelial cell interaction, and this cell pair was not found in any other tissue group. To demonstrate such findings, there are bolded chords for all the healthy tissue chemokine interactions in healthy skin (**Figure 3a**), healthy lung (**Figure 3c**), healthy colon (**Figure 3e**) and healthy kidney (**Figure 3g**). In the disease datasets, we observed both tissue-exclusive and non-exclusive sender-receiver pairs. Tissue specificities for sender-receiver pairs involved in cell-cell communication are also indicated in **Supplemental Table 5** (for the healthy tissues) and **Supplemental Table 6** (for the disease-tissues). The subsequent sections will elaborate on the tissue-specific interactions in more detail.

**Figure 3.**
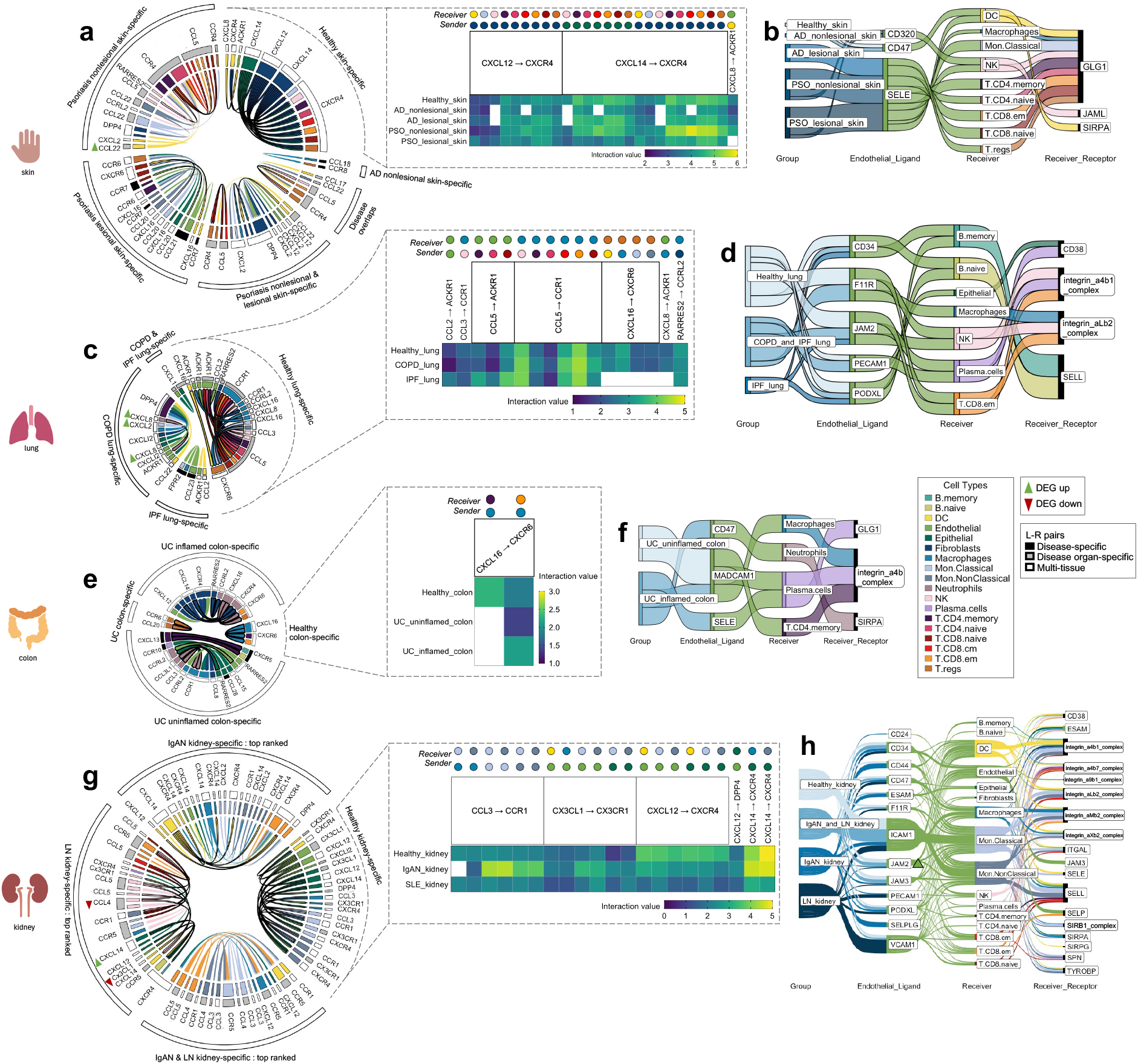
The tissue-specific sender-receiver ligand-receptor cell-cell communications identified with CellphoneDB; segmented by the four different organs (skin, lung, colon, kidney).Heatmaps display healthy tissue chemokine communications (which typically co-occurred in disease with either increased/decreased interaction values, i.e. with directional changes). Chord diagrams display healthy-tissue specific and disease-specific chemokine communications. Sankey diagrams display immune cell extravasation communications. **a)** Healthy skin, atopic dermatitis and psoriasis-specific chemokine-based communication **& b)** extravasation-based communication. **c)** Healthy lung, chronic obstructive pulmonary disease and idiopathic pulmonary fibrosis-specific chemokine-based communication **& d**) extravasation-based communication. **e)** Healthy colon and ulcerative colitis-specific chemokine-based communication **& f)** extravasation-based communication. **g)** Top-ranked healthy kidney, IgA nephropathy and lupus nephritis-specific chemokine-based communication **& h)** extravasation-based communication. *Bolded chords indicate sender_cell-receiver_cell communication that are unique to a particular disease. AD* atopic dermatitis, COPD chronic obstructive pulmonary disease, *DEG* differently expressed gene, *IgAN* IgA nephropathy, *IPF* idiopathic pulmonary fibrosis, *les* lesion, LN lupus nephritis, *nl* nonlesion, *PSO* psoriasis, *UC* ulcerative colitis.

#### 3.3.1 Healthy skin, atopic dermatitis, and psoriasis-specific interactions

There were 26 chemokine cell-cell communications specific to healthy skin, of which 25 involved CXCL12/CXCL14 --> CXCR4, and one CXCL8 --> ACKR1 communication (Figure 3a, blowout panel). In nearly every instance, these communications co-occurred in AD and PSO (**Figure 3a**). Most of the sender cell types in healthy skin were non-immune cell types (Fibroblasts and Epithelial cells), and their receivers were DCs, NKs, and all of the T cell subtypes (**Figure 3a**).

There were 47 chemokine cell-cell communications specific to lesional/nonlesional PSO and nonlesional AD skin, only two of which overlapped across these three disease groups. There were no interactions specific to lesional AD skin. The notable feature differentiating cell-cell communications in diseased skin vs healthy skin was the presence of signals originating in immune cells. Of all the healthy and disease tissues examined, DC/Endothelial/Mon.Classical --> T.CD8.naive communications only occurred in lesional PSO skin. Moreover, the Endothelial --> T.CD8.naive communication involved CCL21 --> CCR7 signaling, an L-R pair that was only detected in lesional PSO skin.

There were nine skin-specific extravasation-based communications (**Figure 3b**). Most of them took place in AD and PSO skin, with only one that occurred in healthy skin, involving Endothelial_CD320 communication to DC_GLG1. From the DEG analysis, there were no differentially expressed genes that were common across the PSO lesional skin datasets used in this study (**Supplemental Figure 1a**).

However, a DEG finding that was found in both datasets of nonlesional PSO skin was a higher transcript abundance of DC_CCL22 (p < 0.01). DC_CCL22 was also involved in nonlesional PSO skin-specific chemokine cell-cell interactions to T.regs and T.CD4.naive cells (**Figure 3a**), meaning that the increased DC expression of CCL22 may be promoting its cell communication to these T cells.

#### 3.3.2 Healthy lung, chronic obstructive pulmonary disease, and idiopathic pulmonary fibrosis-specific interactions

There were 17 chemokine cell-cell communications identified that were specific to healthy lung. Most of these communications were also present in COPD and IPF, with the exception of CXCL16 --> CXCR6 and CXCL8 --> ACKR1 interactions, which were absent in IPF lung (**Figure 3c**). Compared to other tissues, Macrophages commonly served as the receiver cell in the healthy lung, more specifically for CCL3 --> CCR1, RARRES2 --> CCRL2, and multiple communications involving CCL5 --> CCR1. As well, unlike the healthy skin chemokine-based communications, healthy lung did not involve Fibroblasts and Epithelial cells as the senders, but rather myeloid cells and T cells. In contrast, Endothelial and Epithelial cells were the most common senders in COPD and IPF lung-specific cell-cell chemokine communications (**Figure 3c**). Nonetheless, Neutrophil --> Endothelial communication was exclusively present in COPD lung tissue. Intriguingly, communication from the opposite direction, from Endothelial --> Neutrophil for extravasation was absent (**Figure 3d**), raising the possibility of reverse transendothelial migration, a phenomenon previously observed *in vitro* (Buckley et al., 2006).

Across the lung groups, there were seven different extravasation-based communications (**Figure 3d**). Although each of the endothelial ligands present in healthy lung co-occurred in COPD and/or IPF lung, none of the corresponding receivers or receptors matched those observed in healthy lung. This suggests the presence of lung immune cell extravasation mechanisms unique to diseased states that do not occur under constitutive conditions.

Differential expression analysis revealed several DEGs in the COPD lung data, specifically, higher transcript abundances of Macrophage_CXCL2 (p < 0.001), Epithelial_CXCL1 (p < 0.01), and Neutrophil_CXCL8 (p < 0.05) compared to healthy lung (**Supplemental Figure 1b**). COPD lung also displayed elevated expression of Macrophage_CXCL8 (p < 0.001), which was involved in cell-cell communication in both healthy and COPD lung tissues; COPD lung displayed a slightly higher interaction value, possibly attributed to the differential expression (**Figure 3c**).

#### 3.3.3 Healthy colon and ulcerative colitis-specific interactions

In the healthy colon, only two chemokine cell-cell communications were identified. Both involved Macrophage_CXCL16 --> T.CD4+/CD8+ memory cells expressing CXCR6, with only the latter communication co-occurring in both uninflamed and inflamed UC colon (**Figure 3e**). Intriguingly, the communication between Macrophage_CXCL16 --> T.CD4.memory_CXCR6, observed exclusively in healthy colon, stood out as the sole communication in this study not found in the corresponding diseased tissue of that organ.

In UC inflamed and uninflamed colon, a total of 19 chemokine cell-cell communications were identified, with only one being specific to uninflamed UC (**Figure 3e**). In inflamed UC colon, unique sender-receiver pairs involved T.CD4.memory --> B.naïve/B.memory, Endothelial/Fibroblast --> Neutrophils, and Neutrophils --> Macrophages. Interestingly, extravasation-based communication was observed for both Macrophages and Neutrophils in inflamed UC colon (**Figure 3f**). Moreover, the mentioned T.CD4.memory --> B.naïve/B.memory communications involved CXCL13 --> CXCR5, an L-R pair only found in inflamed UC colon. In uninflamed UC colon, a unique sender-receiver pair was Neutrophils --> T.CD8.em/T.regs, as well as T.regs --> T.regs; however, these cells did not seem to be extravasating (**Figure 3f**). There were no genes from the DEG analysis that co-occurred across all the UC datasets. However, cell proportion comparisons unveiled that T.regs, known for their role in anti-inflammation, were actually higher in proportion in inflamed UC colon compared to healthy colon (p < 0.01), a paradoxical finding supported by previous research (Himmel et al., 2012; Lord et al., 2015) (**Supplemental Table 3** as indicated earlier).

#### 3.3.4 Healthy kidney, IgA nephropathy, and lupus nephritis-specific interactions

There were 21 chemokine cell-cell communications specific to healthy kidney, and all except one (Macrophage_CCL3 --> Mon.Classical_CCR1) also co-occurred in IgAN and LN kidney (**Figure 3g**). These interactions revealed that the same cell types were usually involved regardless of the ligand-receptor pair, with various combinations of myeloid/Endothelial/Epithelial cell --> myeloid/DC communication occurring across CCL3 --> CCR1, CX3CL1 --> CX3CR1 and CXCL14 --> CXCR4 L-R pairs. In IgAN kidney, there were 248 cell-cell chemokine cell-cell communications specific to this condition, while for LN kidney, there were 171, making diseased kidney the most communication-rich tissue. There were 48 communications that overlapped between the two diseases; the top 20 interactions with the highest interaction value within each of these three sectors were plotted (**Figure 3g**). IgAN had unique sender-receiver pairs including DCs --> DC/Macrophage/Mon.Classical/Mon.NonClassical/T.CD8.em cells, while LN had unique sender-receiver pairs including Endothelial cells --> NKs, NKs --> T.CD8.em/DC/Mon.NonClassical, T.CD8.naïve --> Mon.NonClassical, T.CD8.cm --> T.CD8.em/Mon.Classical/Mon.NonClassical, and Epithelial --> B.naïve/B.memory (**Figure 3g**). All these receiver cells also exhibited extravasation-based communication in both IgAN and LN kidney (**Figure 3h**).

Interestingly, there was higher transcript abundance of Endothelial_JAM2 in LN kidney (p < 0.05, **Supplemental Figure 1c**), overlapping with several extravasation-based communications occurring in the kidneys of healthy, IgAN and LN conditions (**Figure 3h**). LN kidneys also showed higher proportions of most annotated immune cell types, including B.memory, B.naive, DC, Macrophages, Mon.Classical, Mon.NonClassical, NK, Plasma cells, T.CD4.memory, T.CD4.naive, T.CD8.cm, T.CD8.em, T.CD8.naive and T.regs, and a significantly lower proportion of Epithelial cells than healthy kidney (p < 0.05, **Supplemental Table 3** as indicated earlier), raising the possibility of there being increased cell communication in LN due to the increased amounts of immune cells present.

#### 3.3.5 Healthy and disease tissue common interactions

Seven chemokine cell-cell communications occurred in more than one healthy tissue organ, though none took place in the colon (**Figure 4a**). Importantly, these healthy tissue interactions that co-occurred in multiple healthy tissue types were not found in any disease tissues. Considerably more (58) chemokine cell-cell communications co-occurred in disease, of which the Endothelial_CXCL12 --> T.CD8.em_CXCR4 communication was present in all 10 disease groups examined in this study (**Figure 4b**). These disease multi-tissue interactions did not occur in any healthy tissues.

**Figure 4.**
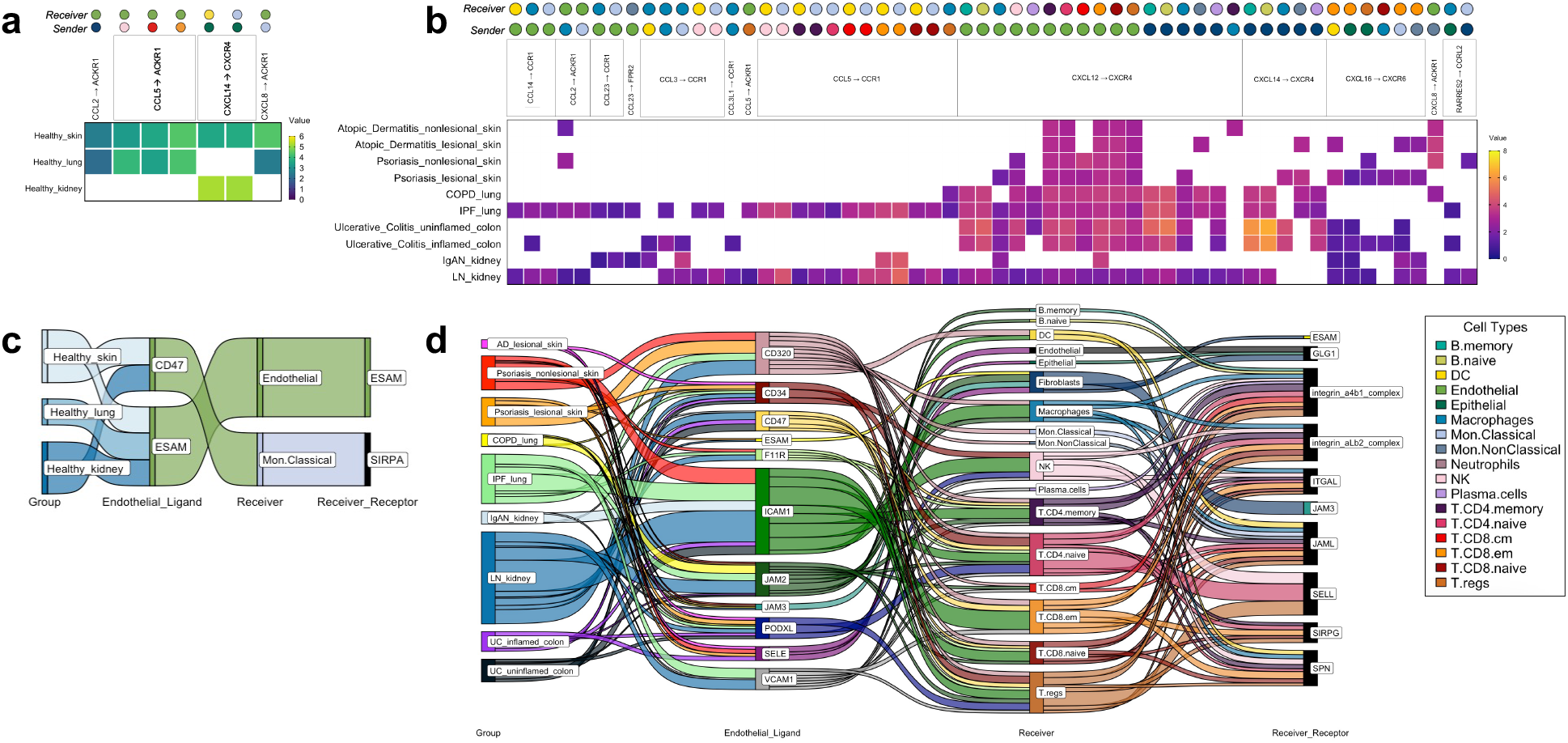
The tissue-nonspecific sender-receiver ligand-receptor cell-cell communications identified with CellphoneDB, with heatmaps for chemokine-based communications and Sankey diagrams for immune cell extravasation-based communications. **a)** Healthy chemokine-based communication that were common across tissues, specifically, healthy skin, healthy lung and healthy kidney, at either increased/decreased interaction strength values. **b)** Disease chemokine-based communication that were common across all 10 disease groups examined. **c)** Healthy tissue extravasation-based communication that were common across three of the tissues examined. **d)** Disease extravasation-based communication that were common across nine of the disease groups examined. *AD* atopic dermatitis, COPD chronic obstructive pulmonary disease, *IgAN* IgA nephropathy, *IPF* idiopathic pulmonary fibrosis, LN lupus nephritis, *PSO* psoriasis, *UC* ulcerative colitis.

There were five common extravasation-based communications occurring across multiple healthy tissues (**Figure 4c**). As with the chemokine-based communications, these involved all the healthy tissues studied except for colon and none of them were found in the disease tissues. In the diseases examined, there were numerous (198) common extravasation-based interactions occurring across lesional AD skin, PSO nonlesional and lesional skin, COPD and IPF lung, IgAN and LN kidney, and uninflamed and inflamed UC colon, and amongst these communications, the receivers spanned almost all of the cells that are annotated by SignacX, apart from Neutrophils (**Figure 4d**).

### 3.4 Experimental investigation

In the IgAN kidney dataset, we found cell-cell communications which can be considered atypical chemokine ligand to receptor bindings that have not yet been investigated in chemotaxis assays. This included: 1) CXCL2 signaling, known as a neutrophil chemoattractant (Li et al., 2016), to T.CD8.em cells via DPP4, also known as CD26 and 2) RARRES2, an adipokine (Tang et al., 2023), signaling to classical and nonclassical monocytes via chemokine receptor CCRL2. First, we measured chemotaxis of pan CD8+ T cells and pan monocytes towards their respective target chemokines in a transwell assay and found that CXCL2 did indeed chemoattract CD8+ T cells; specifically, 1 ng/ml of CXCL2 had a significantly higher migration index compared to the medium-only control (p < 0.05 for both donors, **Supplemental Figure 2a**). CXCL12, a chemokine well-known to chemoattract T cells (Hara et al., 2006), also led to significantly higher migration (p < 0.001 for both donors). However, RARRES2 did not chemoattract monocytes (**Supplemental Figure 2b**), unlike CCL8 (also known as monocyte chemoattractant protein-2 (MCP-2)) which led to significantly higher migration for both donors, particularly at the 0.1 ng/ml concentration (p < 0.01, **Supplemental Figure 2c**), compared to the negative controls.

The data and t-test results for the transwell assays are reported in **Supplemental Table 8** and **Supplemental Table 9**, respectively.

After establishing which of the tested ligands chemoattracted the predicted cell type(s), we asked two follow-up questions. First, we aimed to determine whether receptor expressions would change in the presence of the chemoattracting ligand, and specifically, a) if the expression of DPP4 and CXCR2, the canonical receptor of CXCL2 (Zhang et al., 2017), increases in the presence of CXCL2 post-chemotaxis and b) if CXCR4, the canonical receptor of CXCL12 (Shi et al., 2020), increases when CXCL12 is expressed, and c) if the expression of CD25, an activation marker (Bajnok et al., 2017), increases post- chemotaxis. Second, we tested the involvement of DPP4 in the CXCL2-induced chemoattraction of CD8+ effector memory T cells by knocking out DPP4.

To answer the first question, we conducted the CXCL2 transwell assay with the CD8+ effector memory T cells and then collected the migrated cells for flow cytometry quantification (**Figure 5a**).

**Figure 5.**
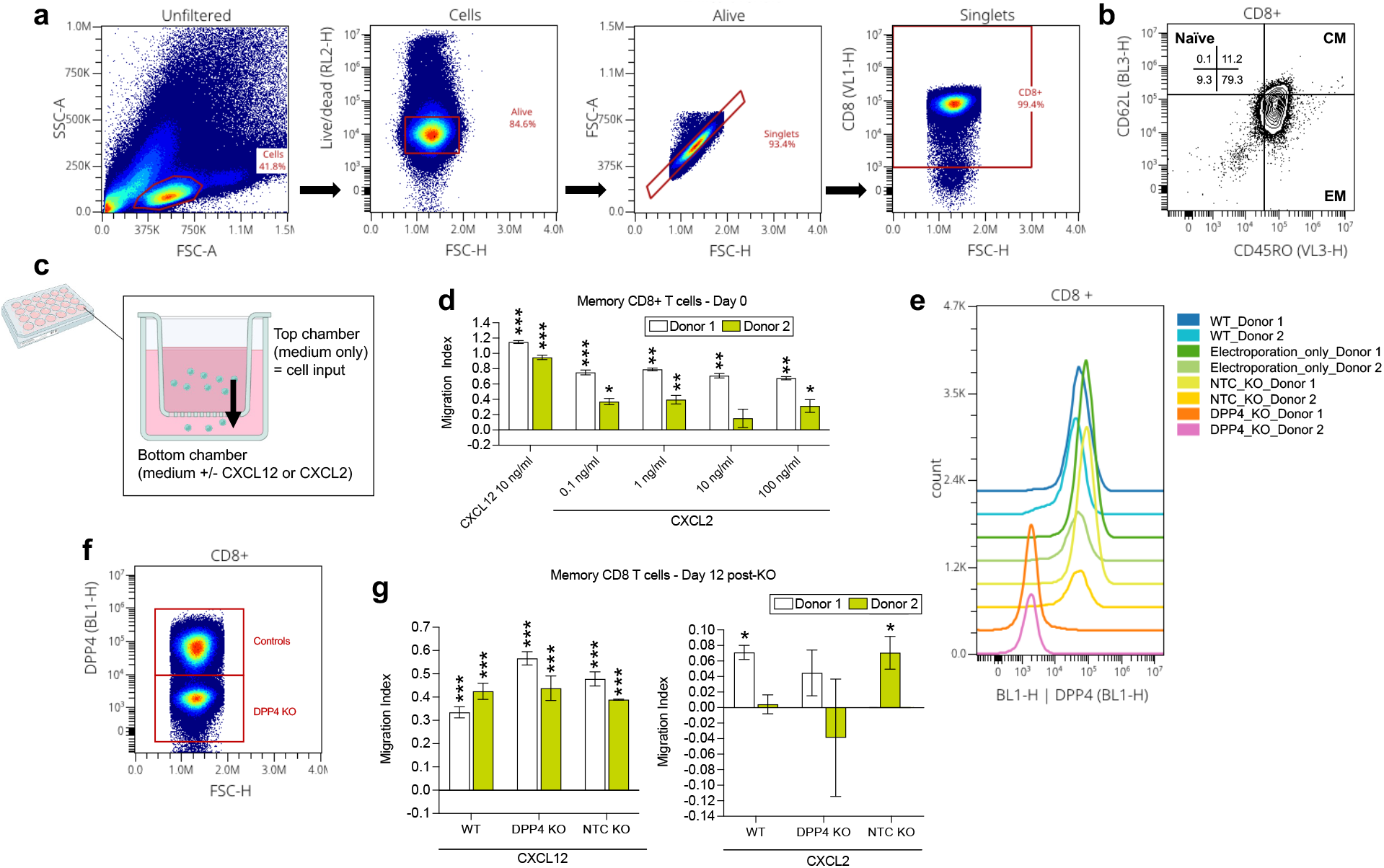
Experimental investigation of the CXCL2 --> CD8 T effector memory cell_DPP4 interaction found in the IgAN kidney cell-cell communication analysis. **a)** Flow cytometry analysis of CD8+ memory T cells. **b)** Identification of CD8+ memory T cells as predominantly effector memory (EM) subtype (CD45RO+ and CD62L-), with a minor fraction of central memory (CM) cells (CD45RO+ and CD62L+). **c)** Cartoon depiction of the chemotaxis assay using the transwell; overtime, cells inputted to the top chamber of the transwell may be chemoattracted to the bottom chamber, similar to how within tissues, a chemotactic gradient chemoattracts cells to a site of inflammation. **d)** CXCL2 transwell migration assay conducted on day 0 revealed chemotaxis of CD8+ memory T cells towards CXCL2 (at all dilutions examined for both donors, with the exception of 10 ng/ml for donor 2), consistent with the meta-analysis findings. **e-f)** In CD8+ memory T cells, reduced DPP4 expression was observed only in the DPP4 KO group, while the wild-type (WT), electroporation only, and non-target control (NTC) KO groups showed no significant alterations on day 11. **g)** Subsequent CXCL2 transwell migration assay conducted with CD8+ memory T cells to assess the impact of DPP4 KO on CXCL2-induced chemotaxis. DPP4 KO cell’s CXCL2-induced migration was not significantly different from the medium-only condition (for both donors), contrasting with wild-type (WT) and NTC cells in donor 1 and 2 respectively, that showed significantly stronger migration towards CXCL2 compared to the medium-only control (p < 0.05). CXCL12 positive control showed similar levels of significantly higher migration compared to the medium-only control (p < 0.05 for both donors) for all groups (KO, WT, NTC). * p < 0.05. ** p < 0.01, *** p < 0.0001 compared to the medium control condition per donor.

Ultimately, there were no significant changes in the levels of expression of DPP4, CXCR2, CXCR4, or CD25. However, we did find that memory CD45RO+ CD8+ T cells, most of which were effector memory cells (CD62L-CD45+; **Figure 5b**), migrated towards CXCL2 (**Figure 5c**). Like the pan CD8+ T cell findings, 1 ng/ml CXCL2 led to significantly higher migration compared to the medium-only condition (p < 0.05 for both donors). Next, we assessed if knocking out DPP4 would affect the ability of the CD8+ memory T cells to migrate to CXCL2 (**Figure 5 d-f**). We determined that DPP4 KO cell’s CXCL2-induced migration was not significantly different from the medium-only condition, contrasting with wild-type (WT) and NTC cells in donor 1 and 2 respectively, that showed significantly higher migration towards CXCL2 compared to the medium-only control (p < 0.05). In contrast, the CXCL12 positive control showed similar levels of significantly higher migration compared to the medium-only control regardless of treatment group (p < 0.001 for WT, KO, NTC for both donors). These findings suggest that the CXCL2-DPP4 axis might indeed play a role in the chemoattraction of CD8+ effector memory T cells.

## 4. DISCUSSION

By integrating data from publicly available scRNA-seq datasets, we created a directory of shared and disease-specific immune cell chemokine and extravasation ligand-receptor (L-R) cell-cell communications occurring in multiple immune-mediated diseases. This analysis was conducted across diverse conditions affecting major organs, AD and PSO skin, COPD and IPF lung, UC colon, and IgAN and LN kidney. We found a varying degree of specificity among interactions, ranging from being specific to a particular diseased tissue to occurring across multiple healthy and diseased organs. Our analysis also delved into associated alterations in cell proportions and DEGs that may be a cause and/or response to enhanced immune cell infiltration. As proliferation and activation marker DEGs were absent in our findings, chemotaxis and immune cell extravasation were likely the primary drivers of the predicted cell-cell communications, rather than cell proliferation within tissue. This work resulted in a novel, *in silico-*derived roadmap of immune cell migration, offering a navigable resource for extracting genes of interest for experimental testing, such as for the exploration of the effects of L-R inhibition on disease pathology. Further, the analytical framework employed for extracting our findings can be applied to future analyses, enabling the incorporation of additional tissues as new datasets and diseases come into focus.

The aim of this study was to shed light on the disease-specificity and tissue-specificity of L-R signals driving immune cell migration in immune-mediated and autoimmune diseases. Considering the L-R pairs found in our analysis, none were exclusive to healthy tissue; all the L-R pairs found in healthy tissue co-occurred in one or more disease tissue for both chemokine and extravasation-based communication (**Figure 2a, b**). A similar pattern was observed when considering all four components of cell-cell communication — the sender cell (i.e. the cell expressing the ligand), its ligand, the receiver cell (i.e. the cell receiving the ligand from the sender, with its receptor), and its receptor. Specifically, we found communications that were specific to particular healthy tissues (heatmaps in **Figure 3**); however, nearly all (65; 98%) these communications co-occurred in their respective diseases, with the exception of one found in healthy colon (Macrophage_CXCL16 --> T.CD4.memory_CXCR6) and those seven that were common across healthy tissues (**Figure 4a**). This suggests a gain of healthy tissue interactions in disease conditions rather than a complete switch to disease-enriched ones. This lack of healthy tissue-specific cell communication aligns with past research suggesting that gene interactions involved in immune pathology often stem from processes inherent to constitutive occurrences (Peterson and Artis, 2014; Ardain et al., 2020).

A notable difference with respect to signaling specificity was observed between chemokine and extravasation-based communication. Among the L-R pairs identified, there was a higher prevalence of exclusivity to disease tissue, termed “disease-enriched”, within the chemokine-based L-R pairs (**Figure 2a**; 72% of all pairs), compared to the extravasation-based L-R pairs (**Figure 2b**; 33% of all pairs). These findings are also consistent with prior research suggesting that unlike chemotactic mechanisms, immune cell extravasation entails more generalized functions across cell types and healthy versus disease states (Garrood et al., 2006; Nguyen and Soulika, 2019).

There was only one case where context-free cell communication occurred, whereby the Endothelial_CXCL12 --> T.CD8.em_CXCR4 interaction was observed across all the diseases examined (**Figure 4b**). Not considering the cell types involved, the CXCL12--> CXCR4 and CXCL14--> CXCR4 signals were the only ones ubiquitously observed in all the diseases examined, as well as in some healthy tissues (**Figure 2a**). This finding coincides with past research that suggests the fundamental role of CXCL12 and CXCL14 in immune cell recruitment throughout the body (García-Cuesta et al., 2019).

Previous research also suggests that CXCL12 and CXCL14 exert opposing effects on CXCR4, with CXCL12 acting as an agonist and CXCL14 as an antagonist (Hara and Tanegashima, 2014). Interestingly, in both healthy and disease skin interactions (**Figure 3c**) and UC inflamed colon (**Figure 3e**), it was predicted that CXCL12 and CXCL14 from fibroblasts communicate towards the same immune cell types (neutrophils in UC, and in skin: DC, Mon.Classical, NK, T.CD4.memory (& naïve), T.CD8.naive(& central memory, effector memory), T.regs). This redundancy in cell communication highlights the complex interplays between chemokines in disease pathology (Turner et al., 2014; Li et al., 2022).

The “disease organ-specific” L-R pairs identified for both chemokine (**Figure 2a**) and extravasation-based communication (**Figure 2b**) underscore the potential presence of signals that are specific to tissue disease states. For example, the CCL5 --> CCR4 interaction was exclusively observed in the context of diseased skin, involving various types of T cells in communication (**Figure 3c**). This tissue specificity in T-cell mediated CCL5 --> CCR4 interactions was not attributable to the composition of niche cell types in the skin, as T cells were also present in lung (**Figure 3a**), colon (**Figure 3e**) and kidney (**Figure 3g**) disease interactions. Moreover, in such scenarios, disease appeared to influence the effectiveness of the L-R signals. For example, the CCL5 --> CCR5 interaction involving NK -->T.regs was shared across AD and psoriatic skin, however, this same L-R pair was also detected across broader categories of T cells as senders and receivers in PSO skin. The variability in immune cell migration patterns elicited by a single L-R signal across different diseases underscores the complexity of immune-mediated disease pathology. Understanding how these signals adapt in diverse disease contexts could inform the design of more effective treatments.

Another major observation in our study is the abundance of kidney-specific L-R pairs (**Figure 2a, b**). In the case of extravasation-based communication, this contrasts with the relatively limited L-R pairs utilized by the diseased skin endothelium (**Figure 3d**; SELE/CD47/CD320 to GLG1/SRPA/JAML). The intricate signaling network observed in kidney, with 13 endothelial ligands and 18 receiver cell receptors involved in extravasation (**Figure 3h**) likely reflects the unique architecture of kidney tissue, which requires fine control of immune infiltration (Suárez-Fueyo et al., 2017). Another finding that reflects the influence of tissue architecture on immune cell communication is the CXCL13 --> CXCR5 memory T cell to B cell interaction that was specific to inflamed ulcerative colitis colon (**Figure 3e**), aligning with the presence of secondary lymphoid structures within the colon (Mörbe et al., 2021). Thus, anatomical features are likely involved in determining tissue-specificity in immune cell communication and migration (Weisberg et al., 2021; Correia, 2023).

While tissue-specificity may play a strong role in determining immune cell communication in disease, this may also be due to shared disease etiologies, such as a reliance on type 2-associated immune responses in both IgAN and LN (Ebihara et al., 2001; Ko et al., 2022). An additional limitation to our study is that just four tissues were considered. The addition of more immune disease datasets, such as those from the brain, bones, or muscle, may have led to different findings regarding tissue-specificity and commonalities. Our findings may also be influenced by variations in the numbers of patients and cells amongst the scRNA-seq datasets included in our analysis. However, our approach of extracting findings that co-occur within specific tissue groups, such as healthy kidney or COPD lung, serves to not only yield more conservative conclusions, but also acts as a means to address this limitation. Our cell-cell communication predictions, as well as cell proportion and DEG findings are not based on protein expressions, but rather mRNA levels, which often, but not always, represent protein abundance (Liu et al., 2016). Despite that, CellphoneDB, compared to other cell-cell interaction tools, has shown better performance in consistency with spatial tendency (Liu et al., 2022). Moreover, our mRNA-based predictions of the expression of certain ligands or receptors in particular cells are supported by immunohistochemical studies of diseased human tissue (even though such studies are limited), including elevated epithelial CCL20 in psoriasis ((Homey et al., 2000); **Figure 3a**) and endothelial CX3CL1 in IgAN kidney ((Cox et al., 2012); **Figure 3g**).

Another limitation of *in silico* predictions of cell-cell communication involves the necessity of experimental validation or support from existing literature. In this study, we adopted an approach where chemotaxis assays were employed using predicted ligands as chemoattractants, and knock-outs of predicted receptors within the anticipated ’receiver’ cell types served to validate the predicted L-R interactions. Our findings revealed that CCL8, but not RARRES2, chemoattracted monocytes as predicted in the IgAN kidney dataset (**Supplemental Figure 2a, 2b**). While an *in vitro* invalidated result challenges the accuracy of *in silico* predictions, it could also suggest that RARRES2 signals to monocytes, but the presence of multiple chemokines might be necessary to facilitate its tissue penetration (Luster et al., 2005). Also through the chemotaxis assay, we observed that the CXCL2 --> T.CD8.em_DPP4 interaction identified in the IgAN kidney dataset may indeed be genuine, particularly evidenced by the reduced CXCL2-induced migration upon DPP4 knockout (**Figure 5**). Such experimental approaches hold promise for future validation of novel cell-cell communications, thereby enhancing the reliability of such findings in identifying cell signals implicated in disease pathogenesis.

In conclusion, we found that chemotactic cues vary in their specificity, with some potentially restricted to particular disease tissue. Our work contributes to the understanding of disease-specific and non-specific ligand-receptor cell communication, which can support biomarker discovery and precision medicine efforts aimed at increasing treatment efficacy. Further, it can bolster targeting strategies involving chemokine signaling pathways, potentially resulting in tissue-specific approaches aimed at inhibiting immune cell chemotaxis to reduce inflammation.

## AUTHOR CONTRIBUTIONS

Conceptualization: MFR, MV, VS, TKM, AK, Data curation: MFR, AK, Formal analysis: MFR, Funding acquisition: VS, EdR, TKM, Investigation: MFR, Methodology: MFR, MV, EM, TKM, AK, Project administration: MFR, Resources: VS, EdR, TKM, MV, EM, NN, Software: AK, MFR, NN, Supervision: MFR, AK, MV, TKM, VS, NN, Validation: MFR, AK, Visualization: MFR, Writing – original draft: MFR, VS, AK, Writing – review & editing: MFR, AK, MV, EM, VS, NN.

## ACKNOWLEDGEMENTS

We extend our gratitude to the researchers and developers of the multiple scRNA-seq datasets, algorithms and packages used for our meta-analysis, as their contributions were integral to the production of this study. The healthy kidney and lupus nephritis-related work was supported by the Accelerating Medicines Partnership® Rheumatoid Arthritis and Systemic Lupus Erythematosus (AMP® RA/SLE) Program. AMP® is a public-private partnership (AbbVie Inc., Arthritis Foundation, Bristol-Myers Squibb Company, Foundation for the National Institutes of Health, GlaxoSmithKline, Janssen Research and Development, LLC, Lupus Foundation of America, Lupus Research Alliance, Merck & Co., Inc., National Institute of Allergy and Infectious Diseases, National Institute of Arthritis and Musculoskeletal and Skin Diseases, Pfizer Inc., Rheumatology Research Foundation, Sanofi and Takeda Pharmaceuticals International, Inc.) created to develop new ways of identifying and validating promising biological targets for diagnostics and drug development Funding was provided through grants from the National Institutes of Health (UH2-AR067676, UH2-AR067677, UH2-AR067679, UH2-AR067681, UH2-AR067685, UH2- AR067688, UH2-AR067689, UH2-AR067690, UH2-AR067691, UH2-AR067694, and UM2- AR067678).

We would also like to acknowledge Frank O. Nestle, MD for his contributions to the conceptualization of this study.

## DATA AVAILABILITY

This work relied on publicly available datasets listed in Supplemental Table 2 with the exception of the AMP_LN dataset, which was obtained from the ARK portal (arkportal.synapse.org). To ensure full replicability, the parameters used for processing the datasets on CellBridge are listed in Supplemental Table 2 and the html summaries of the outputs are available on https://zenodo.org/records/12754198. Outputs of all the analyses are available in the specific supplemental tables indicated throughout the manuscript.

---

**Supplemental Figure 1.**
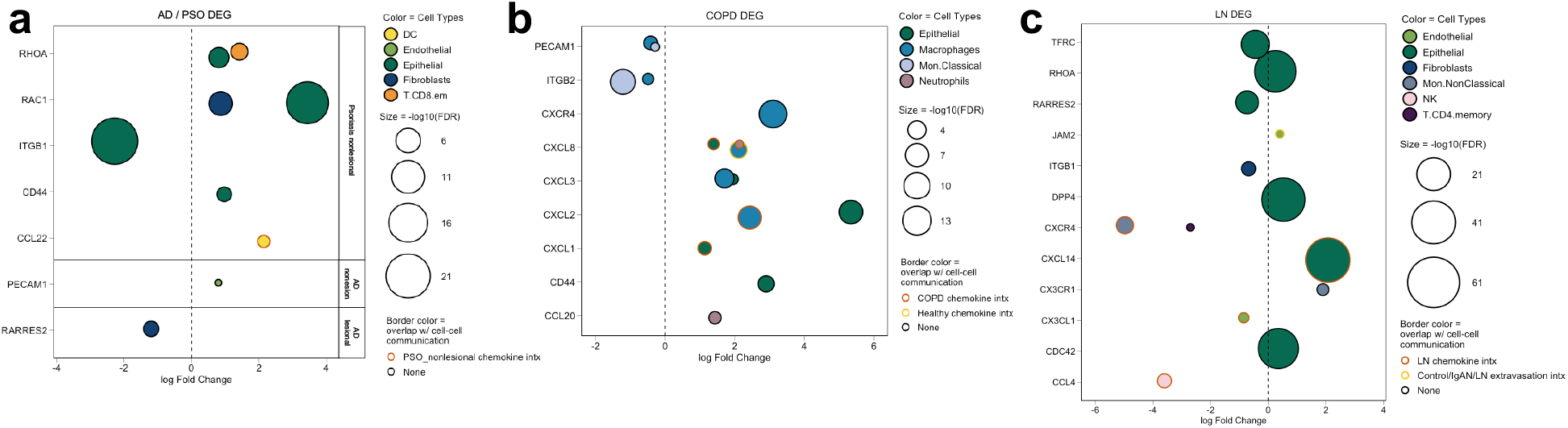
Chemokine and immune cell extravasation differentially expressed genes (DEGs) that occurred in: **a)** Atopic dermatitis (AD) and psoriasis (PSO) skin, **b)** Chronic obstructive pulmonary disease (COPD) lung, **c)** Lupus nephritis (LN) kidney.

**Supplemental Figure 2.**
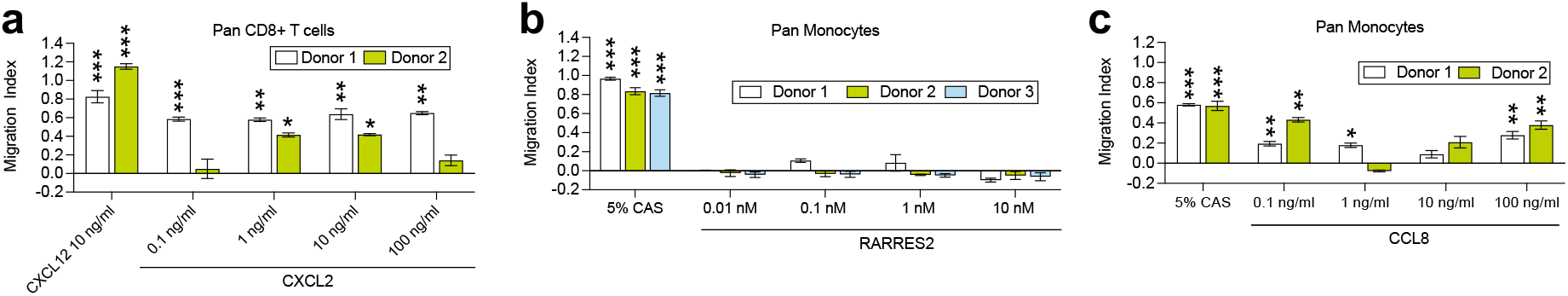
Experimental investigation of ligand – immune cell interactions in transwell migration assays. a) Pan CD8 T cells migrated towards CXCL2 (at all dilutions for donor 1, and at 1 ng/ml and 10 ng/ml for donor 2), **b)** pan monocytes did not migrate towards RARRES2, **c)** pan monocytes migrated towards CCL8 (at 0.1 ng/ml, 1 ng/ml and 100 ng/ml for donor 1, and at 0.1 ng/ml and 100 ng/ml for donor 2).* p < 0.05. ** p < 0.01, *** p < 0.0001 compared to the medium control condition per donor.

